# Ambient temperature, clutch size, and daylight are the main drivers of incubation behavior in two neotropical swallows breeding 8,000 km apart

**DOI:** 10.1101/2025.08.27.672727

**Authors:** David Diez-Méndez, Marcela Liljeström, Caren B. Cooper, Daniel R. Ardia, David Winkler

**Affiliations:** Faculty of Science, University of South Bohemia, České Budějovice, Czechia; Department of Animal Ecology, Netherlands Institute of Ecology (NIOO-KNAW), Wageningen, The Netherlands; Institute of Entomology, Biology Centre of the Czech Academy of Sciences (BC CAS), České Budějovice, Czechia; Centro Austral de Investigaciones Científicas (CADIC), CONICET, Ushuaia, Argentina; College of Natural Resources, North Carolina State University, Raleigh, NC, USA; Department of Biology, Franklin and Marshall College, Lancaster, PA, USA; SABER Consulting, Monterey, CA, USA

**Keywords:** Tachycineta, latitudinal effects, breeding behaviour, incubation constancy, local adaptation

## Abstract

The central dilemma in incubation behaviour is how to allocate time between warming up the clutch and self-preservation activities outside the nest. It is assumed that incubating birds follow a classical framework in which the environmental temperature, the main variable influencing incubation behaviour, and bout duration show a non-linear relationship. In this context, incubating birds would maximize bout duration for most of the temperature range in which they breed. To determine whether this theoretical framework applies to species breeding at different latitudes under extremely different environmental conditions, we collected incubation data from two distant but breeding populations of *Tachycineta* swallows. *T. leucopyga* (Chillean swallow) breeds in Ushuaia, Argentina, and *T. meyeni* (mangrove swallow) breeds in Hill Bank, Belize. These populations do not overlap in terms of ambient temperature; they are exposed to different daylight conditions and the latitudinal differences influence life-history traits such as clutch size. *Tachycineta* swallows showed similar incubation consistency over 24 hours (∼75% of nest attendance) but followed different behavioral strategies to achieve this consistency. Daylight duration limited incubation activity in mangrove swallows, but longer nocturnal incubation compensated for shorter days. Larger clutches only took their toll on the Chilean swallows, by shortening their incubation bouts and lengthening their foraging bouts, which affected the length of their daily activity. Overall, both swallow species showed incubation patterns adapted to the local conditions. They responded differently to increasing temperatures, but both species reduced their incubation effort whenever possible. Our results contrast with previous work where maximizing incubation effort was the rule. In our populations, females reduced their incubation effort as soon as weather conditions improved (i.e., increasing temperatures), changing the paradigm of incubation behaviour.

**Lay summary:**

- In species where only the male or female incubates, they need to find a balance between the time they spend incubating in the nest and the time they spend foraging out of the nest.
- The ambient temperature is the most important factor influencing the duration of both behaviours, and birds are thought to respond similarly to the same range of temperatures.
- We studied the incubation behaviour of two species of swallows breeding in Belize and Argentina, respectively.
- Both species incubated for a similar length of time in 24 hours, but each followed a different strategy depending on clutch size and daylight duration.
- Their incubation behaviour was also related to ambient temperature, but differed between populations, suggesting adaptations to local conditions.
- We cannot expect different species to show similar responses to ambient temperature under different conditions.

## Introduction

The balance between incubating bird’s physiology requirements and the thermal requirements of the clutch is the central dilemma during avian incubation (White and Kinney 1974, Conway and Martin 2000). In uniparental incubating species with intermittent diurnal behavior, the focal bird needs to optimize the duration of self-maintenance activities, e.g., foraging and preening, against clutch cooling (Haftorn 1981, Deeming 2002). If the nest temperature falls below the physiological zero temperature (PZT, 26-28°C) during the absence of the incubating parent (off-bouts), embryo development may be slowed down or even suspended (Webb 1987, Haftorn 1988, Olson et al. 2006, 2008). When the incubating parent returns to the nest (on-bouts), it needs to provide a steadily and narrow temperature range (37-40.5°C) (Nord and Nilsson 2011, DuRant et al. 2013), to avoid developmental impairment (Drent 1975, Webb 1987, Olson et al. 2008), additional metabolic costs (Hepp et al. 2006, Olson et al. 2006, Ardia et al. 2010, Nord and Nilsson 2016), loss of hatchability (Hepp et al. 2006, Macdonald et al. 2014), and reduced survival rates in the short and long-term (Berntsen and Bech 2016, Ospina et al. 2018). On and off-bouts are therefore the basic constituents of an individual’s incubation behavior, and their interplay is assessed as the nest attentiveness, i.e., the proportion of time a bird is incubating the clutch, which can be calculated hourly, daily, or for the full incubation period (Diez-Méndez et al. 2021a).

Early studies have shown that off-bout and on-bout duration depends on ambient temperature, in a non-linear relationship (White and Kinney 1974, Conway and Martin 2000). Conway and Martin (2000) proposed a theoretical framework in which incubating birds are assumed to maximize their bout duration following increasing ambient temperature up to the PZT. This is followed by a stable phase in which there is no correlation between ambient temperature and bout duration, up to 40.5°C (upper thermal limit). Since the on-bouts are usually many times longer than the off-bouts and change their length at a higher rate, incubating birds that maximize the duration of the on-bouts ultimately increase their incubation effort. However, numerous studies addressing avian incubation behaviour at different latitudes have shown that the timing of on-bouts and off-bouts does not conform to Conway and Martin’s framework: either the stable stage is shifted from the 26°C – 40.5°C range (Kovařík et al. 2009), there is no observable stable stage for on- and/or off-bouts (Amininasab et al. 2016, Diez-Méndez et al. 2021a) or bout duration is negatively correlated with increasing temperatures (Morton and Pereyra 1985, Camfield and Martin 2009, Macdonald et al. 2014, Coe et al. 2015, Schöll et al. 2019, Diez-Méndez and Cunningham 2024). These a priori divergent patterns of incubation behavior have been described in different species but also between populations of the same species thriving under different environmental conditions (Nord and Cooper 2020, Lundblad and Conway 2021, Diez-Méndez et al. 2021a). Instead of maximizing incubation effort (Conway and Martin 2000), these studies suggest that there is an inflection point in the interplay between off- and on-bouts where time allocation shifts from increasing incubation effort to increasing off-nest self-care; although this inflection point may be based on local ambient temperatures rather than fixed values (Diez-Méndez et al. 2021a).

Despite local adaptations, it is possible to observe common latitudinal patterns of incubation behaviour that are explained by life-history theory (Saether 1988, Ricklefs 2000). At low latitudes, birds lay smaller clutches, show lower nest attendance, and longer incubation periods (Cooper et al. 2005, Hille and Cooper 2015, Jin et al. 2025, but see Austin et al. 2019). Smaller clutches exhibit higher heat dissipation and appear to be favored in warmer habitats/lower latitudes (egg-viability hypothesis, Veiga 1992, Cooper et al. 2005, Boulton and Cassey 2012), while larger clutches show higher thermal inertia and appear to be favored in higher latitudes/cooler habitats (egg-cooling hypothesis, Reid et al. 2002, Cooper et al. 2005, Boulton and Cassey 2012). The evolutionary causes for the lower nest attendance and the longer incubation periods at low latitudes (Robinson et al. 2008) are still controversial (Ricklefs et al. 2017, Martin et al. 2018). On the one hand, several studies support the hypothesis that mainly intrinsic physiological programs and not external factors such as temperature or survival play a role in embryonic developmental schedules (Tieleman et al. 2004, Robinson et al. 2008, 2014; Ricklefs and Brawn 2013, Austin et al. 2019). On the other hand, the main role of incubation temperature and adult bird survival on incubation periods and nest attentiveness is also widely supported (Martin 2002, Martin et al. 2007, 2013, 2018). Another external factor, the duration of daylight, may also contribute to shortening incubation periods by accelerating embryo growth programming at high latitudes and prolonging them at lower latitudes (Cooper et al. 2011, Austin et al. 2014). From a physiological perspective, incubating birds at high temperatures are constrained more by energy requirements than by daylight for their daily activities, whereas at lower latitudes the shorter daylight duration is likely to be the constraint (Shaw and Cresswell 2014). This is already observed at mid-latitudes, where incubating birds end their active day up to three hours before sunset (Diez-Méndez et al. 2021a). At low latitudes, the expected lower nest attendance can otherwise be compensated by a longer night time with continuous incubation behavior (Shaw and Cresswell 2014, Sofaer et al. 2020, Nord and Cooper 2020).

Despite the numerous studies addressing incubation behavior, few of them compare in detail the same or closely related species across different latitudes, and of these, studies of incubation behavior are limited to a few latitudinal degrees and the Northern Hemisphere (Ardia et al. 2006a, Rohwer and Purcell 2019, Sofaer et al. 2020, Nord and Cooper 2020, Lundblad and Conway 2021). In this study, we investigated the incubation behaviour of two *Tachycineta* species that breed more than 8,000 km apart: *Tachycineta albilinea* (mangrove swallows) breed in the Northern Hemisphere but at low latitude in the tropics, while *Tachycineta leucopyga* (Chilean swallows) breed at high latitude in the Southern Hemisphere. In our study case, *T. albilinea*, breeding in Hill Bank (Belize), shows larger clutches and shorter incubation periods than *T. leucopyga*, which breeds in a harsh habitat in Ushuaia (Argentina). Our aim was to compare the incubation behavior of these two species, in which ambient temperatures do not overlap during the breeding season, and to analyze whether their bouts and nest attendance follow an overarching pattern driven by ambient temperature or whether they are adapted to local conditions. We calculated on- and off-bout duration to determine hourly and daily nest attendance, 24h incubation, and active day duration. We hypothesized that both species share common patterns that are likely to be widespread in incubating passerines but follow different strategies (Austin et al. 2019): an increase on incubation effort towards the end of the incubation period (Cooper and Voss 2013), an effect of clutch size (Nord and Nilsson 2012), an daytime effect on bout duration and nest attendance (Conway and Martin 2000, Diez-Méndez et al. 2021a), and finally a non-linear temperature effect on bout duration (Conway and Martin 2000). However, we expect that local conditions determine bout duration and nest attendance, causing different dynamics between on- and off-bouts since the ambient temperature scenarios and energetic requirements are expected to be different.

## Methods

### Study species and breeding sites

We have studied two species of Tachycineta swallows that breed in nest-boxes: the Chilean swallow (*T. leucopyga*) in Ushuaia, Argentina, and the mangrove swallow (*T. albilinea*) in Hill Bank, Belize. Chilean swallows have the southernmost breeding range within their genus and are austral migrants that stay in the Ushuaia area from October to March. In contrast, mangrove swallows are non-migratory and breed from March to July in a tropical area.

Chilean swallow females that breed in Ushuaia lay between 2 and 5 eggs with a mode of 4 eggs (Liljesthröm et al. 2012), which is comparable to *Tachycineta* swallows at low latitudes and smaller compared to *Tachycineta* swallows elsewhere at high latitudes (Ardia et al. 2006a). For example, mangrove swallows (*Tachycineta albilinea*) in Panama (9° 9’ N, 79° 50’ W) laid 3-5 eggs with a mean of 4 eggs, while white-rumped swallows (*Tachycineta leucorrhoa*) in Buenos Aires Province (35° 34’ S, 58° 01’ W) laid 4-6 eggs with a mode of 5 eggs, and tree swallows (*Tachycineta bicolor*) in Fairbanks, Alaska, USA (64° 49’ N, 147° 52’ W) laid a mean (sd) of 5.7 (0.66) eggs (Ardia et al. 2006a).

Argentina: During the 2008-2009 breeding season, we studied incubation behavior of female Chilean swallows breeding in nest-boxes at two sites near the city of Ushuaia, Tierra del Fuego, Argentina. Site A (54° 44’ S, 68° 12’ W) was located 11 km northeast of Ushuaia, and Site B (54° 53’ S, 67° 20’ W) was located 85 km east of Ushuaia and had a total of 88 and 62 nest-boxes, respectively. The study area is part of the Andean-Patagonian woodland biogeographic region. The monthly average ambient temperature (sd) varies between 9.2 °C (4.3 °C) in December and 10.4 °C (4.2 °C) in January. Daily temperatures range from -0.5 °C to 24 °C. Daylength, measured by civil twilight, was between 16.5 and 19.3 hours during the breeding season.

Belize: During the 2009 breeding season, we studied incubation behavior of female mangrove swallows breeding in a nest-box population in Hill Bank, Belize (17.60 N, 88.70 W). The monthly mean ambient temperature (sd) ranged between 29.5 °C (2.75 °C) in May to 28.5 °C (2.89 °C) in June. Daily temperatures range from 23 °C to 37 °C. Daylength by civil twilight varied between 13.7 and 14.7 hours during the breeding season.

During the incubation periods of the 2008-2009 breeding season for *T. leucopyga* and 2009 for *T. albilinea*, we recorded their incubation rhythms by placing temperature recorders (Thermochron iButton dataloggers, accuracy ± 0.5 °C, model DS1922L-F5, Maxim Integrated, USA) in 18 and 10 nests, respectively. The nests were selected according to their laying date in order to include nests throughout the entire breeding season. We placed 2 iButtons in each nest box on the day of the third egg was laid. One iButton was placed inside the nest cup to measure the temperature in the environment directly next to the eggs. The second iButton was placed on the inside of one of the side walls of the nest box to measure the ambient temperature inside the nest-box. The data loggers recorded temperature at 2-minute intervals and were tested during the 2007-08 breeding season without affecting adults’ behavior (no nest abandonment was recorded, unpublished data). The dataloggers did not provide an exact temperature of the eggs during incubation, but they do produce time series with fluctuations consistent with the incubation behavior of uniparental, intermittent incubators (Joyce et al. 2001, Cooper and Mills 2005). A drop in temperature indicates when a female leaves the nest, and a subsequent increase in temperature indicates her return to the nest. For all nests, we also collected data on clutch size and apparent incubation duration (the day on which the last egg of the clutch was laid was considered as incubation day 1).

### Incubation data analysis

We used the program Rhythm (Cooper and Mills 2005) to select incubation recesses from the temperature recordings. Whenever the temperature in the nest cup dropped by more than 2 °C and lasted more than 2 min, we considered that observation as a recess. We then used Raven Pro 1.6 (Cornell Laboratory of Ornithology) to visually inspect the time series selected by Rhythm, and we manually selected any recess that had not been automatically selected.

From this data, we determined the start time and duration of each off-bout and on-bout. Where possible, we also determined when the first off-bout started in the morning and the last on-bout ended in the evening, so that the active day could be assessed. We calculated the hourly nest attentiveness (total number of minutes a female was on the nest per hour) and the daily nest attentiveness (total number of minutes a female was on the nest during her active day). We also calculated 24h incubation behavior assuming that females incubated continuously during the passive day (mainly at night, Pendlebury and Bryant 2005).

We assigned ambient temperature to an off-bout or on-bout by selecting the ambient temperature at the onset of the focal bout. We also calculated the hourly mean temperature for the hourly nest attentiveness and the daily mean temperature (temperature during daylight) for the daily nest attentiveness. When assessing the first off-bout in the morning, we calculated the mean ambient temperature of the ending night. For the assessment of the last on-bout in the evening, we used the aforementioned daily mean temperature. For simplicity, the calendar date was calculated as 1 = 1 April in Belize and 1 = 1 November in Ushuaia.

### Data analysis

To evaluate the relationship between ambient temperature and incubation rhythms, we used linear (LMMs) and generalized mixed-effects models (GLMMs) separately for Argentina and Belize. Mixed models account for the non-independence of data in repeated measurements (in this case, nests were measured several times during the day or in different days) (Pinheiro and Bates 2000). We built two models to assess bout duration: 1) off-bout duration (min), and 2) on-bout duration (min). We built an hour-level model: total time on nest (min/60, hourly nest attentiveness). We modeled three response variables at the daily level: 1) number of off-bouts, 2) total time on nest (min)/active day (min) (daily nest attentiveness), and 3) duration of active day (min). To further investigate the duration of the active day, we built two additional models that analyzed 1) the start of the first daily off-bout in relation to dawn and 2) the end of the last daily on-bout in relation to dusk.

We used R software (R Core Team 2023) to build the models and calculate their estimates. In the bout models we chose a gamma distribution with a log link because of the long right tail the data showed (glmmTMB package, Brooks et al. 2017). Clutch size, incubation day, ambient temperature (quadratic term, *poly-*function), hour of the day (quadratic term, *poly-*function) and calendar date were the fixed-effect variables. Nest ID was a random intercept variable, and incubation day was also nested within Nest ID for a random slope variable, to account for daily variability in slopes. We built two models per bout type, one containing the interaction *Temperature x Hour of the day* and another one without the interaction. We compared them to select the model that better fit our data using a likelihood rate test.

For the hour-level model we chose a gaussian distribution. Clutch size, incubation day, ambient temperature (quadratic term, poly-function), hour of the day (quadratic term, poly-function) and calendar date were the fixed-effect variables. Nest ID was a random intercept variable, and incubation day was also nested within Nest ID for a random slope variable to account for daily variability in slopes. We built two models, one containing the interaction *Temperature x Hour of the day* and another one without the interaction. We compared them to select the model that better fit our data.

For the day-level models we also chose a gaussian distribution. Clutch size, incubation day, ambient temperature (quadratic term, poly-function), and calendar date were the fixed-effect variables. Nest ID was a random intercept variable. Specifically, we substituted calendar date for active day duration (min) in daily nest attentiveness models. We also substituted calendar date for daylight duration (min) in the active day models.

For the daily first off-bout and last on-bout models we also chose a gaussian distribution. Clutch size, calendar date, incubation day, and ambient temperature (quadratic term, poly-function) were the fixed-effect variables. Nest ID was a random intercept variable.

Figures, model estimates and 95% CI were created with *gglplot2* package (Wickham 2016) and *Effects* package (Fox and Hong 2009). All fixed variables were scaled. Data is presented as mean ± SD unless stated otherwise.

## Results

### Incubation rhythms

We collected data on incubation behavior for 28 nests. Each nest was monitored on average for 6.5 days (median: 5.5 days, range 1-16 days). In Ushuaia, females started incubating between November 26 and January 16 and mean ambient temperature measured inside the nest-box during the incubation period was 9.5 ± 4.27°C, while in Hill Bank, females started incubating between May 17 and June 8 and mean ambient temperature was 28.9°C ± 2.88°C (Fig. 1).

**Figure 1.**
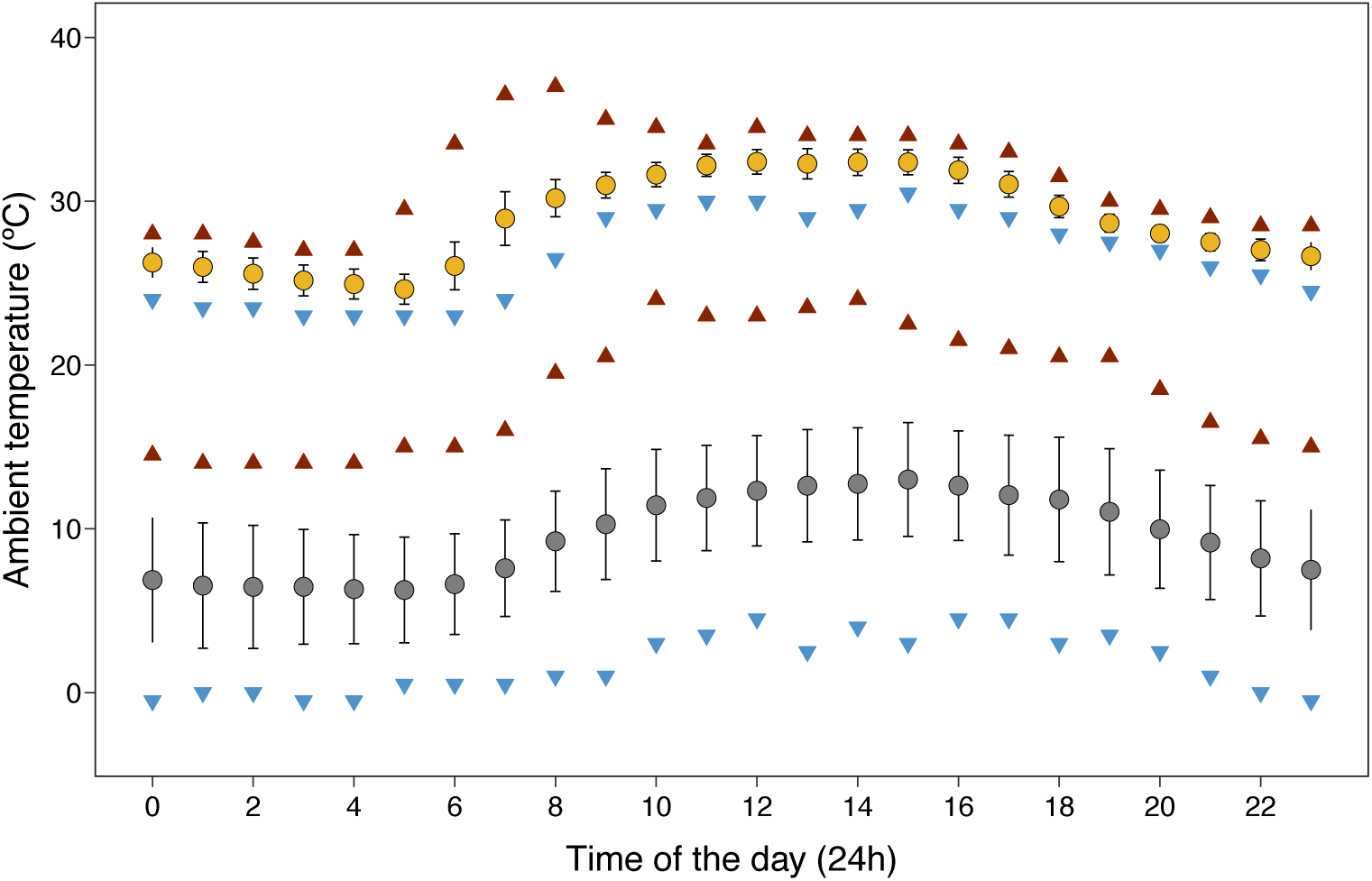
Mean hourly ambient temperature ± SD for Ushuaia, Argentina (grey), and Hill Bank, Belize (yellow) populations. Blue and red triangles show hourly minimum and maximum ambient temperatures respectively.

Chilean swallows laid smaller clutches than mangrove swallows (3.7 ± 0.59 vs. 4.3 ± 0.68, same range for both populations = 3-5 eggs). Chilean swallows left the nest 57 ± 16.9 (range = 22-108) times per day, similar to mangrove swallows at Hill Bank (57 ± 11.1, range = 40-85). Females at both sites spent a median of 6 minutes for self-caring activities out of the nest (i.e., off-bouts duration: Ushuaia: 6 ± 3.7 min, range = 2-46 min; Hill Bank: 7 ± 4 min, range = 2-60 min). Chilean swallows spent a median of 8 minutes in incubating bouts (11 ± 10.8 min, range = 2 – 56 min), whereas mangrove swallows spent a median of 6 minutes (7 ± 4.8 min, range = 2 -260 min) Daily nest attentiveness of Chilean swallows was 63.7 ± 8.03 % and 51.3 ± 5.1 % for mangrove swallows. Chilean swallows not only incubated more intensely but also showed longer active days (940 ± 96.1 min, range = 688-1188 min) compared to Hill Bank females (792 ± 22.7 min, range = 692-810 min), although Hill Bank females made the best use of daylight and were active 95% of the time, in contrast to Ushuaian females (82%). As a result, females at Ushuaia incubated on the nest for ∼10h (599 ± 91.2 min, range = 372 -809 min) during their active day, whereas females at Hill Bank incubated for ∼7h (406 ± 44.5 min, range 330-504 min). Despite the differences in nest attentiveness during the active day, when the active and inactive day, i.e., 24h incubation behavior, were combined females incubated for similar length of time each day: Chilean swallows 76.2% ± 5.93 (1,097 ± 85.4 min, range = 920-1,284 min), Hill Bank 73.3% ± 2.85 (1,056 ± 41.1 min, range =979-1,138 min) (Supp. Mat. Fig. S1). Despite similar 24h incubation times, females in Ushuaia took longer to hatch their clutches than did females at Hill Bank (16.5 ± 0.72 days vs. 14.8 ± 0.79 days, ranging between 15-18 and 14-16 days respectively). Assuming these values of 24h incubation for the duration of the entire incubation period, Chilean swallows needed to incubate their clutches on average ∼40 h longer than mangrove swallows did to hatch them.

### Variation in off-bout duration

Chilean swallows adapted off-bout duration (7,132 bouts, 18 nests) to the time of day and the ambient temperature at that time (Table 1, Figure 2a). In the morning, females shortened their off-bouts as ambient temperature increased, whereas during the rest of the day the relationship between ambient temperature and off-bout duration was subtle. Off-bout duration was also related to the day of incubation: The closer the hatching date was, the shorter the off-bouts were (Table 1).

**Figure 2.**
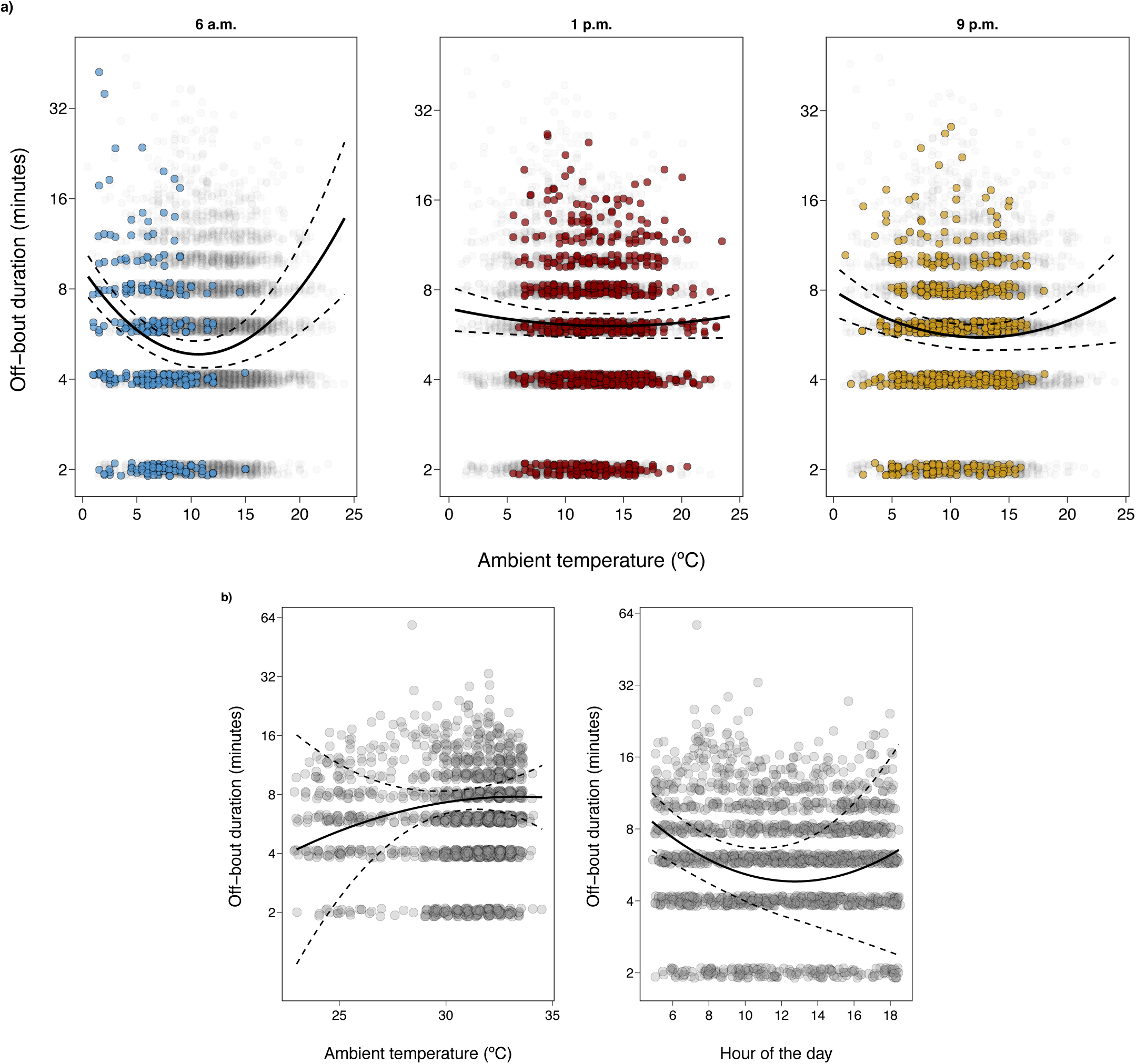
Off-bout duration (log scale) for Chilean swallows (a) and mangrove swallows (b). a) shows the interaction between ambient temperature and hour of the day at three different times. Grey circles show the raw data for off-bout duration. Black continuous and dashed lines show the model estimate and 95%CI respectively. Colored circles show off-bouts that happened from one hour before to one hour after the chosen time.

**Table 1.** Model estimates for the effect of clutch size, incubation day, ambient temperature, hour of the date and calendar date on the **off-bout duration** in **Chilean swallows**. *Nest ID* is a random intercept variable and *Incubation day* is nested as a random slope variable within Nest ID. Significant results are highlighted in bold.

| Fixed effects | Estimate ± SE | z - value | P |
| --- | --- | --- | --- |
| Intercept | 1.78 ± 0.049 | 36.34 |  |
| Clutch size | -0.03 ± 0.056 | -0.58 | 0.563 |
| **Incubation day** | **-0.07 ± 0.030** | **-2.21** | **0.027** |
| Temperature | -0.76 ± 0.812 | -0.93 | 0.351 |
| **Temperature<sup>2</sup>** | **2.81 ± 0.744** | **3.77** | **< 0.001** |
| **Hour of the day** | **1.50 ± 0.651** | **2.31** | **0.021** |
| **Hour of the day<sup>2</sup>** | **-3.45 ± 0.725** | **-4.76** | **< 0.001** |
| Date | 0.08 ± 0.044 | 1.92 | 0.055 |
| Temperature * Hour | -100.08 ± 59.409 | -1.68 | 0.092 |
| **Temperature<sup>2</sup> \* Hour** | **-106.15 ± 50.336** | **-2.11** | **0.035** |
| Temperature * Hour<sup>2</sup> | 52.63 ± 68.791 | 0.76 | 0.444 |
| **Temperature<sup>2</sup> \* Hour<sup>2</sup>** | **170.98 ± 48.490** | **3.53** | **< 0.001** |
| Random effects | Variance ± SD |  |  |
| Nest | 0.034 ± 0.186 | **R<sup>2</sup> fixed effects** | 0.08 |
| Incubation day | 0.009 ± 0.093 | **R<sup>2</sup> model** | 0.22 |

Mangrove swallows shortened off-bouts (1,902 bouts, 10 nests) towards the end of the day (Table 2, Figure 2b). They also tended to lengthen their off-bouts as temperatures increased, but the effect was unclear. As with the Chilean swallows, the closer to hatching day, the shorter the off-bouts were. Larger clutches were also associated with longer off-bouts (Table 2).

**Table 2.** Model estimates for the effect of clutch size, incubation day, ambient temperature, hour of the date and calendar date on the **off-bout duration** in **mangrove swallows**. *Nest ID* is a random intercept variable and *Incubation day* is nested as a random slope variable within Nest ID. Significant results are highlighted in bold.

| Fixed effects | Estimate ± SE | z - value | P |
| --- | --- | --- | --- |
| Intercept | 1.90 ± 0.055 | 34.39 |  |
| **Clutch size** | **0.09 ± 0.028** | **3.2** | **0.001** |
| **Incubation day** | **-0.09 ± 0.039** | **-2.36** | **0.018** |
| Temperature | 6.88 ± 3.846 | 1.79 | 0.074 |
| Temperature<sup>2</sup> | -2.29 ± 2.091 | -1.09 | 0.274 |
| **Hour of the day** | **-3.69 ± 1.738** | **-2.13** | **0.033** |
| **Hour of the day<sup>2</sup>** | **3.47 ± 1.100** | **3.16** | **0.002** |
| Date | -0.04 ± 0.030 | -1.43 | 0.153 |
| Temperature * Hour | 88.35 ± 139.419 | 0.63 | 0.526 |
| Temperature<sup>2</sup> * Hour | 75.56 ± 62.365 | 0.95 | 0.343 |
| Temperature * Hour<sup>2</sup> | -60.85 ± 62.365 | -0.98 | 0.329 |
| Temperature<sup>2</sup> * Hour<sup>2</sup> | 30.94 ± 42.853 | 0.72 | 0.470 |
| Random effects | Variance ± SD |  |  |
| Nest | 0.005 ± 0.074 | **R<sup>2</sup> fixed effects** | 0.09 |
| Incubation day | 0.010 ± 0.100 | **R<sup>2</sup> model** | 0.14 |

### Variation in on-bout duration

Chilean swallows adjusted the duration of on-bouts (6,933 bouts, 18 nests) depending on the time of day and ambient temperature at that time (Table 3, Figure 3a). Overall, incubating females shortened their on-bouts as temperatures rose and lengthened them throughout the day. Larger clutches were associated with shorter on-bouts. We also found a trend towards longer on-bouts later in the season (Table 3).

**Figure 3.**
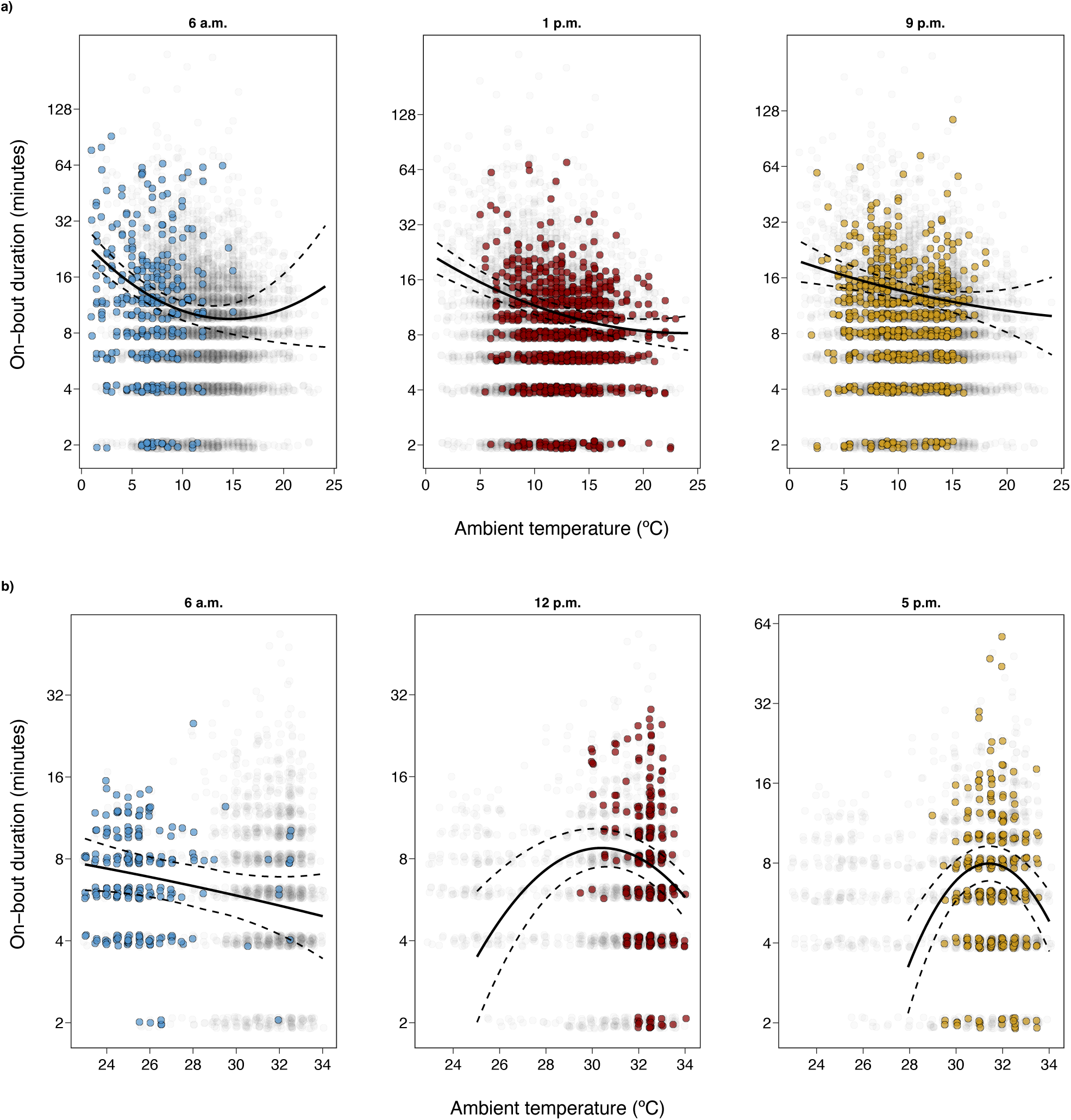
On-bout duration (log scale) for Chilean swallows (a) and mangrove swallows (b). Figures shows the interaction between ambient temperature and hour of the day at three different times. Grey circles show the raw data for off-bout duration. Black continuous and dashed lines show the model estimate and 95%CI respectively. Colored circles show off-bouts that happened from one hour before to one hour after the chosen time.

**Table 3.** Model estimates for the effect of clutch size, incubation day, ambient temperature, hour of the date and calendar date on the **on-bout duration** in **Chilean swallows**. *Nest ID* is a random intercept variable and *Incubation day* is nested as a random slope variable within Nest ID. Significant results are highlighted in bold.

| Fixed effects | Estimate ± SE | z - value | P |
| --- | --- | --- | --- |
| Intercept | 2.44 ± 0.057 | 42.92 |  |
| **Clutch size** | **-0.13 ± 0.044** | **-2.99** | **0.003** |
| Incubation day | -0.08 ± 0.063 | -1.26 | 0.208 |
| **Temperature** | **-12.24 ± 1.026** | **-11.94** | **<0.001** |
| **Temperature<sup>2</sup>** | **2.97 ± 0.936** | **3.17** | **0.002** |
| **Hour of the day** | **5.80 ± 0.826** | **7.03** | **<0.001** |
| **Hour of the day<sup>2</sup>** | **1.87 ± 0.913** | **2.05** | **0.040** |
| Date | 0.07 ± 0.039 | 1.89 | 0.059 |
| Temperature * Hour | 18.04 ± 75.255 | 0.24 | 0.811 |
| **Temperature<sup>2</sup> \* Hour** | **-142.93 ± 66.33** | **-2.15** | **0.031** |
| Temperature * Hour<sup>2</sup> | 131.20 ± 84.610 | 1.55 | 0.121 |
| Temperature<sup>2</sup> * Hour<sup>2</sup> | 44.35 ± 63.952 | 0.69 | 0.488 |
| Random effects | Variance ± SD |  |  |
| Nest | 0.038 ± 0.196 | **R<sup>2</sup> fixed effects** | 0.13 |
| Incubation day | 0.055 ± 0.234 | **R<sup>2</sup> model** | 0.31 |

Mangrove swallows also adjusted on-bout duration (1,830 bouts, 10 nests) depending on time of day and ambient temperature at that time (Table 3, Figure 3b). At the lower and upper end of the daytime temperatures (morning and midday, respectively), on-bouts shortened with increasing temperature. In the evening, the quadratic effect showed a maximum on-bout duration at ∼31-32°C. In contrast to the Chilean swallows, larger clutches were associated with longer on-bouts. We also found a trend towards longer on-bouts later in the season (Table 4).

**Table 4.** Model estimates for the effect of clutch size, incubation day, ambient temperature, hour of the date and calendar date on the **on-bout duration** in **mangrove swallows**. *Nest ID* is a random intercept variable and *Incubation day* is nested as a random slope variable within Nest ID. Significant results are highlighted in bold.

| Fixed effects | Estimate ± SE | z - value | P |
| --- | --- | --- | --- |
| Intercept | 1.83 ± 0.0587 | 21.10 |  |
| **Clutch size** | **0.13 ± 0.024** | **5.36** | **<0.001** |
| Incubation day | -0.13 ± 0.100 | -1.29 | 0.199 |
| **Temperature** | **-13.41 ± 3.731** | **3.60** | **<0.001** |
| **Temperature<sup>2</sup>** | **-10.43 ± 2.044** | **-5.10** | **<0.001** |
| Hour of the day | -2.80 ± 1.600 | -1.75 | 0.081 |
| **Hour of the day<sup>2</sup>** | **-5.69 ± 1.013** | **-5.61** | **<0.001** |
| Date | 0.06 ± 0.035 | 1.78 | 0.075 |
| **Temperature \* Hour** | **621.35 ± 130.93** | **4.75** | **<0.001** |
| **Temperature<sup>2</sup> \* Hour** | **-314.41 ± 74.56** | **-4.22** | **<0.001** |
| **Temperature \* Hour<sup>2</sup>** | **166.62 ± 60.46** | **2.76** | **0.006** |
| Temperature<sup>2</sup> * Hour<sup>2</sup> | -46.78 ± 40.209 | -1.16 | 0.245 |
| Random effects | Variance ± SD |  |  |
| Nest | 0.047 ± 0.217 | **R<sup>2</sup> fixed effects** | 0.10 |
| Incubation day | 0.091 ± 0.302 | **R<sup>2</sup> model** | 0.44 |

### Hourly nest attentiveness

Chilean swallows (2,014 hours, 18 nests) decreased their incubation time as temperatures increased (Table 5). Females incubated less at midday, while evening incubation efforts were higher than in the morning (Table 5). Mangrove swallows (466 hours, 10 nests) also reduced their incubating effort as temperatures increased, but more markedly after midday (Fig. 4). They also increased their incubation efforts throughout the day (Table 6). Calendar date was associated with an increase in hourly nest attentiveness in Hill Bank swallows.

**Figure 4.**
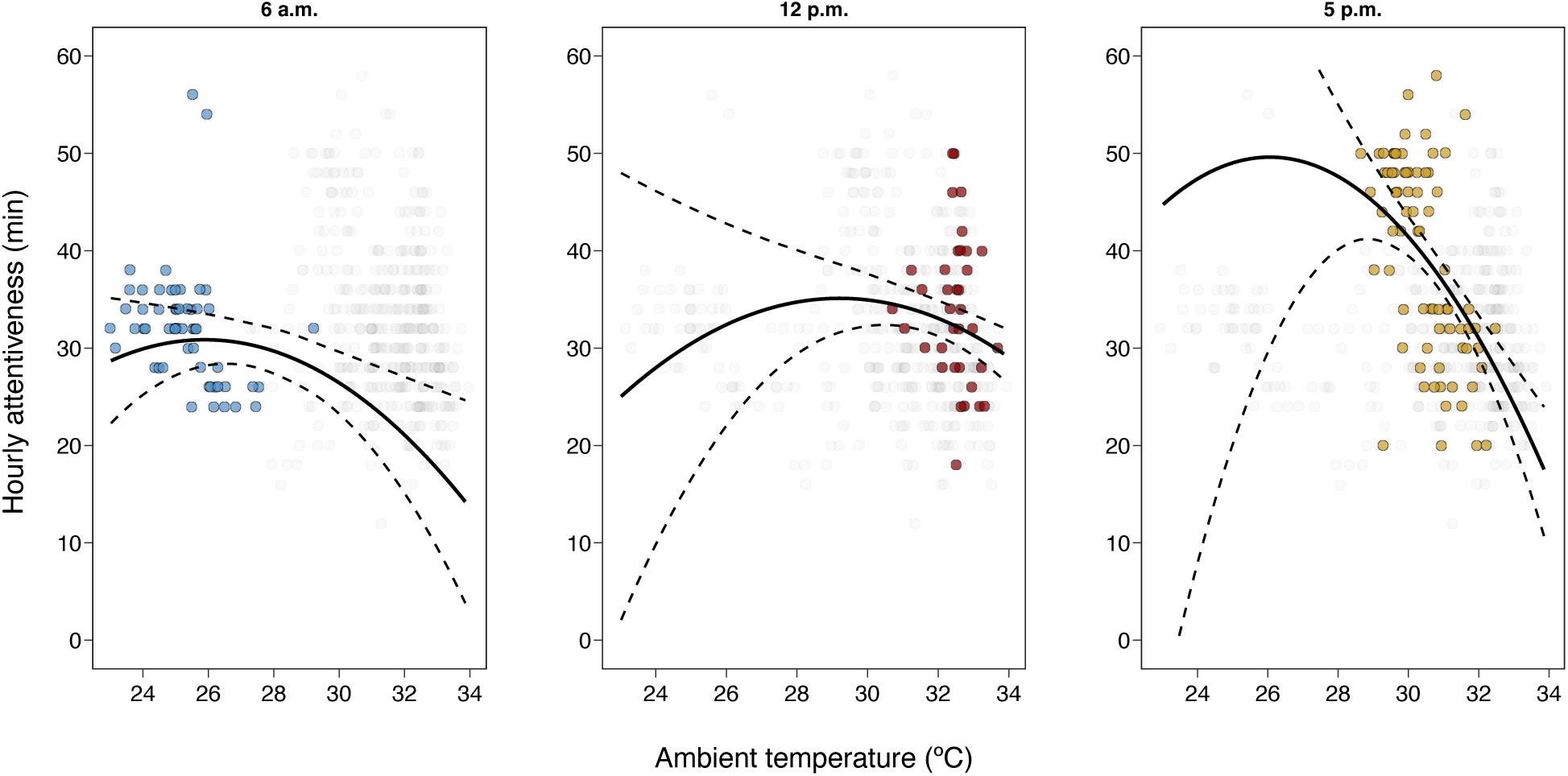
Hourly nest attentiveness for mangrove swallows. Figures shows the interaction between ambient temperature and hour of the day at three different times. Grey circles show the raw data for off-bout duration. Black continuous and dashed lines show the model estimate and 95%CI respectively. Colored circles show off-bouts that happened before 7 a.m, between 11 am and 1 pm, and after 4 pm, respectively.

**Table 5.** Model estimates for the effect of clutch size, incubation day, ambient temperature, hour of the date and calendar date on the **hourly nest attentiveness** in **Chilean swallows**. *Nest ID* is a random intercept variable and *Incubation day* is nested as a random slope variable within Nest ID. Significant results are highlighted in bold.

| Fixed effects | Estimate ± SE | z - value | P |
| --- | --- | --- | --- |
| Intercept | 40.27 ± 1.119 | 35.99 | |
| Clutch size | -1.20 ± 0.958 | -1.26 | 0.208 |
| Incubation day | -0.12 ± 0.935 | -0.13 | 0.900 |
| **Temperature** | **-75.70 ± 9.180** | **-8.25** | **<0.001** |
| Temperature<sup>2</sup> | 7.49 ± 7.279 | 1.03 | 0.303 |
| **Hour of the day** | **16.94 ± 8.139** | **2.37** | **0.018** |
| **Hour of the day<sup>2</sup>** | **122.70 ± 8.139** | **15.08** | **<0.001** |
| Date | -0.079 ± 0.868 | -0.09 | 0.928 |
| Random effects | Variance ± SD | | |
| Nest | 15.70 ± 3.962 | **R<sup>2</sup> fixed effects** | 0.18 |
| Incubation day | 10.07 ± 3.174 | **R<sup>2</sup> model** | 0.47 |

**Table 6.** Model estimates for the effect of clutch size, incubation day, ambient temperature, hour of the date and calendar date on the **hourly nest attentiveness** in **mangrove swallows**. *Nest ID* is a random intercept variable and *Incubation day* is nested as a random slope variable within Nest ID. Significant results are highlighted in bold.

| Fixed effects | Estimate ± SE | z - value | P |
| --- | --- | --- | --- |
| Intercept | 31.17 ± 1.245 | 25.05 | |
| Clutch size | 0.85 ± 0.502 | 1.70 | 0.090 |
| Incubation day | -0.93 ± 0.837 | -1.12 | 0.265 |
| Temperature | 64.23 ± 43.209 | -1.49 | 0.137 |
| Temperature<sup>2</sup> | -47.06 ± 25.71 | -1.83 | 0.067 |
| **Hour of the day** | **89.31 ± 21.671** | **4.12** | **<0.001** |
| **Hour of the day<sup>2</sup>** | **-34.90 ± 17.49** | **-2.00** | **0.046** |
| **Date** | **1.40 ± 0.708** | **1.98** | **0.048** |
| Temperature * Hour | -705.34 ± 719.09 | -0.98 | 0.327 |
| Temperature<sup>2</sup> * Hour | -303.54 ± 466.99 | -0.65 | 0.516 |
| **Temperature \* Hour<sup>2</sup>** | **-1385.79 ± 406.207** | **-3.41** | **0.001** |
| Temperature<sup>2</sup> * Hour<sup>2</sup> | -209.88 ± 278.832 | -0.75 | 0.452 |
| Random effects | Variance ± SD | | |
| Nest | 2.517 ± 1.586 | **R<sup>2</sup> fixed effects** | 0.30 |
| Incubation day | 4.455 ± 2.111 | **R<sup>2</sup> model** | 0.41 |

### Daily nest attentiveness and off-bouts

We were able to record 71 full days (16 nests) of Chilean swallows to calculate nest attentiveness during their active day. The sum of incubating minutes was related to the duration of the active day (Table 7). We also found a trend towards higher nest attentiveness closer to the hatching date (Table 7). Data from the mangrove swallows included 17 full days (9 nests). Females at Hill Bank also extended their incubation time with longer active days (Table 8). In addition, an increase in daily ambient temperature allowed incubating females to reduce their incubation effort after peaking at 30°C, although this relationship was weak (Fig. 5). The later in the season, the higher the daily nest attentiveness.

**Figure 5.**
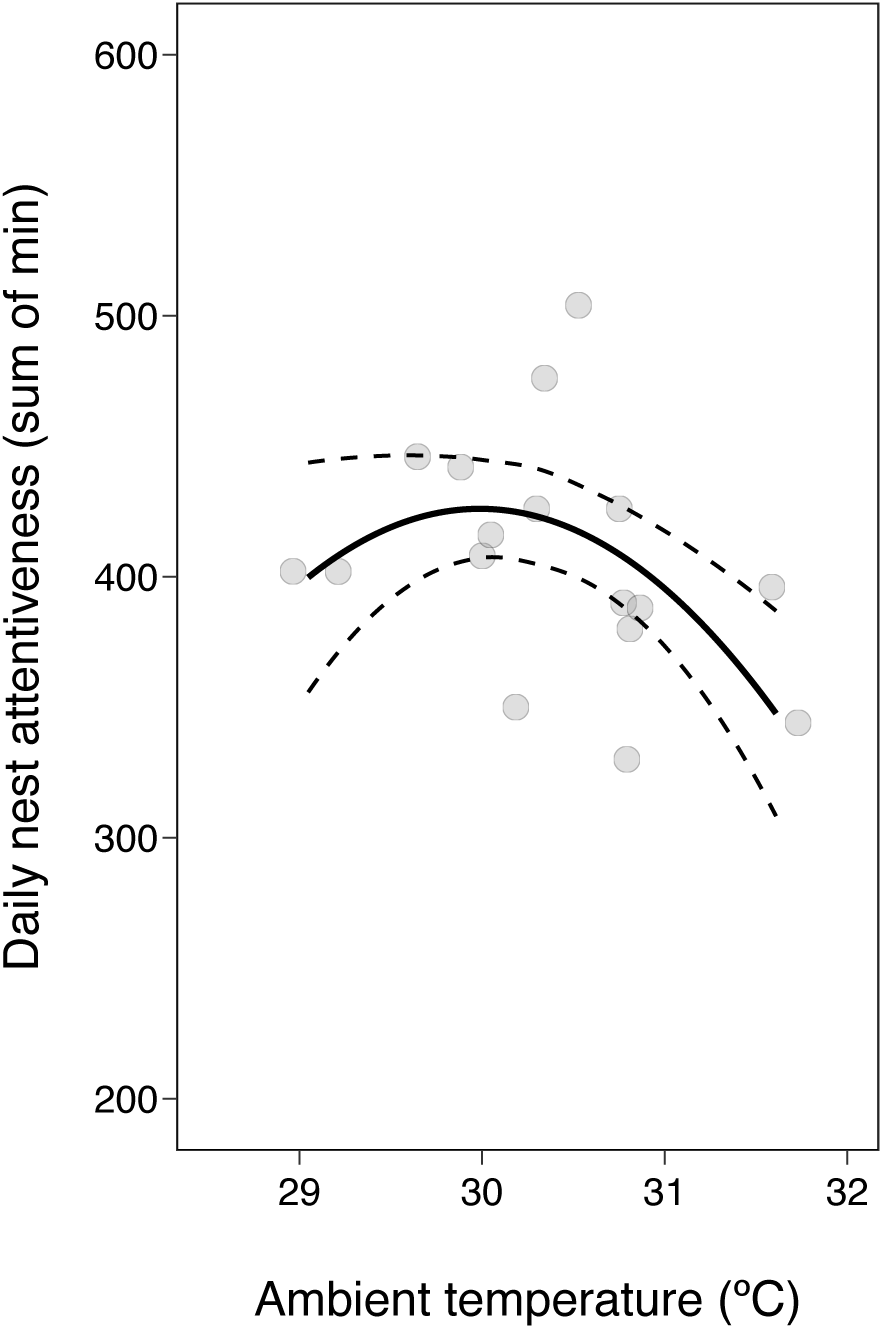
Daily nest attentiveness for mangrove swallows. Grey circles show the raw data for daily incubation (sum of minutes). Black continuous and dashed lines show the model estimate and 95%CI respectively.

**Table 7.**
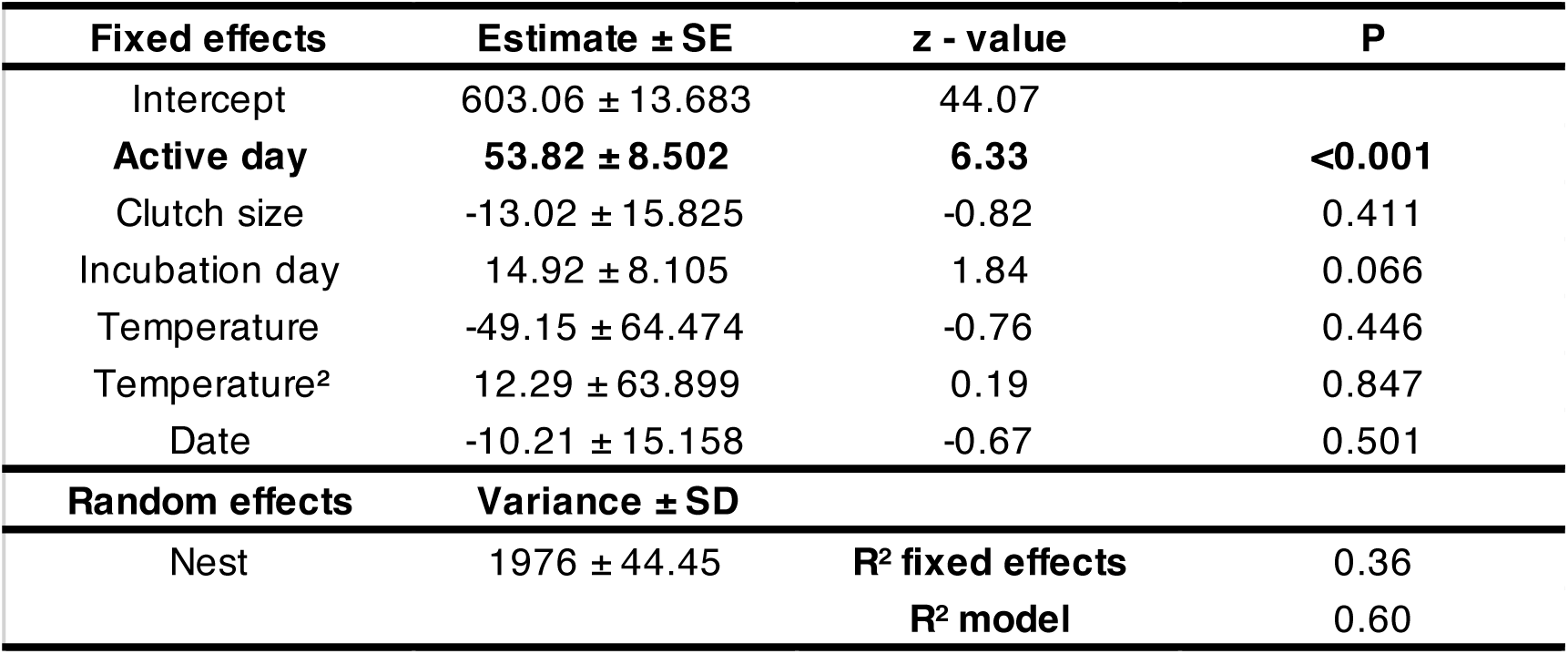
Model estimates for the effect of active day, clutch size, incubation day, ambient temperature, and calendar date on the **daily nest attentiveness** in **Chilean swallows**. *Nest ID* is a random intercept variable. Significant results are highlighted in bold.

| Fixed effects | Estimate ± SE | z - value | P |
| --- | --- | --- | --- |
| Intercept | 603.06 ± 13.683 | 44.07 |  |
| **Active day** | **53.82 ± 8.502** | **6.33** | **<0.001** |
| Clutch size | -13.02 ± 15.825 | -0.82 | 0.411 |
| Incubation day | 14.92 ± 8.105 | 1.84 | 0.066 |
| Temperature | -49.15 ± 64.474 | -0.76 | 0.446 |
| Temperature<sup>2</sup> | 12.29 ± 63.899 | 0.19 | 0.847 |
| Date | -10.21 ± 15.158 | -0.67 | 0.501 |
| Random effects | Variance ± SD |  |  |
| Nest | 1976 ± 44.45 | **R<sup>2</sup> fixed effects** | 0.36 |
|  |  | **R<sup>2</sup> model** | 0.60 |

**Table 8.** Model estimates for the effect of active day, clutch size, incubation day, ambient temperature, and calendar date on the **daily nest attentiveness** in **mangrove swallows**. *Nest ID* is a random intercept variable. Significant results are highlighted in bold.

| Fixed effects | Estimate ± SE | z - value | P |
| --- | --- | --- | --- |
| Intercept | 407.41 ± 6.348 | 64.18 |  |
| **Active day** | **18.45 ± 7.441** | **2.48** | **0.013** |
| Clutch size | -5.31 ± 7.249 | -0.73 | 0.464 |
| Incubation day | 6.45 ± 8.137 | 0.79 | 0.428 |
| **Temperature** | **-56.57 ± 28.346** | **-2** | **0.046** |
| **Temperature<sup>2</sup>** | **-73.35 ± 30.133** | **-2.43** | **0.015** |
| **Date** | **18.13 ± 7.313** | **2.48** | **0.013** |
| Random effects | Variance ± SD |  |  |
| Nest | 0.00 ± 0.006 | **R<sup>2</sup> fixed effects** | 0.65 |
|  |  | **R<sup>2</sup> model** | 0.65 |

In relation to daily nest attentiveness, we investigated the variables that might influence the number of daily off-bouts. For Chilean swallows, clutch size was strongly correlated with the number of off-bouts: the larger the clutch, the greater the number of off-bouts (Table 9). The females also left the nest more frequently on days with average ambient temperatures than on colder or warmer days (Fig. 6). Mangrove swallows tended to leave the nest less frequently when they laid larger clutches, but their daily number of off-bouts was strongly related to the incubation day: more off-bouts when the hatching date was closer (Table 10).

**Figure 6.**
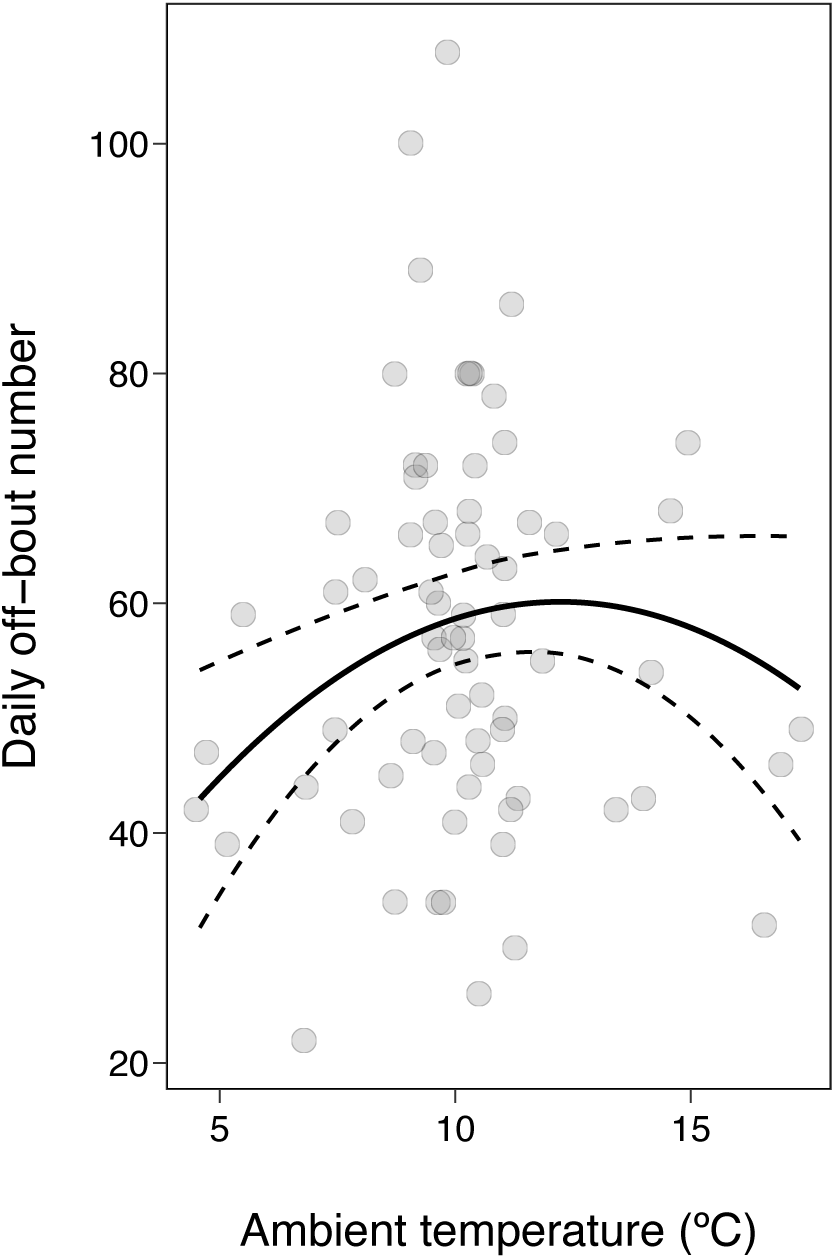
Daily number of off-bouts for Chilean swallows depending on ambient temperature. Grey circles show the raw data for daily incubation (sum of minutes). Black continuous and dashed lines show the model estimate and 95%CI respectively.

**Table 9.**
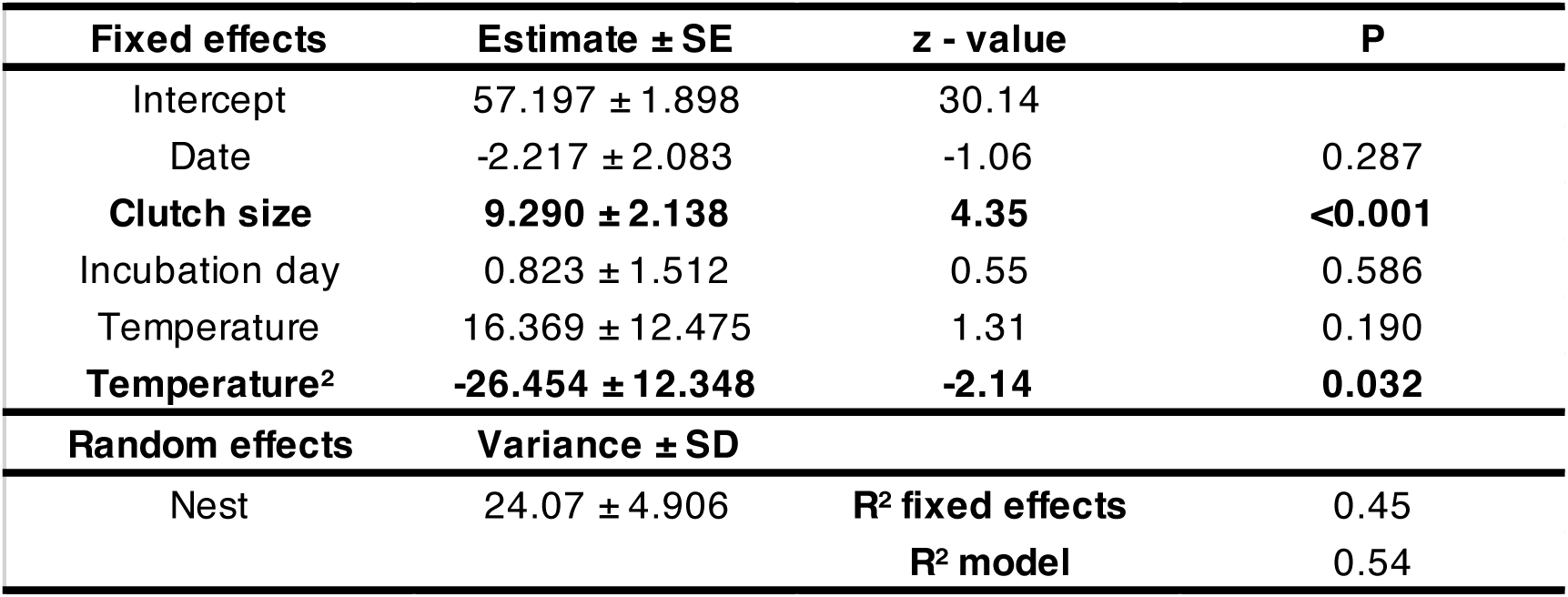
Model estimates for the effect of calendar date, clutch size, incubation day, and ambient temperature on the **daily number of off-bouts** in **Chilean swallows**. *Nest ID* is a random intercept variable. Significant results are highlighted in bold.

| Fixed effects | Estimate ± SE | z - value | P |
| --- | --- | --- | --- |
| Intercept | 57.197 ± 1.898 | 30.14 |  |
| Date | -2.217 ± 2.083 | -1.06 | 0.287 |
| **Clutch size** | **9.290 ± 2.138** | **4.35** | **<0.001** |
| Incubation day | 0.823 ± 1.512 | 0.55 | 0.586 |
| Temperature | 16.369 ± 12.475 | 1.31 | 0.190 |
| **Temperature<sup>2</sup>** | **-26.454 ± 12.348** | **-2.14** | **0.032** |
| Random effects | Variance ± SD |  |  |
| Nest | 24.07 ± 4.906 | **R<sup>2</sup> fixed effects** | 0.45 |
|  |  | **R<sup>2</sup> model** | 0.54 |

**Table 10.**
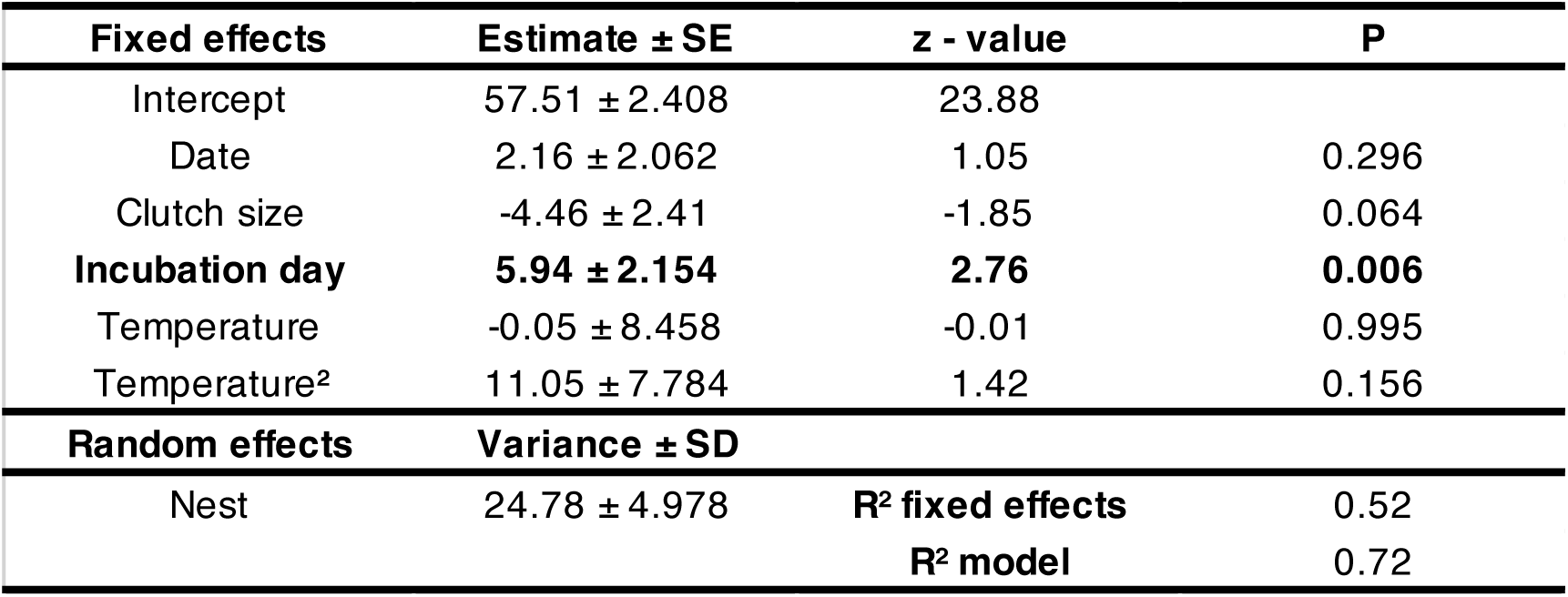
Model estimates for the effect of calendar date, clutch size, incubation day, and ambient temperature on the **daily number of off-bouts** in **mangrove swallows**. *Nest ID* is a random intercept variable. Significant results are highlighted in bold.

### Active day

We found no clear effect of the selected variables on the duration of the active day (71 days, 16 nests) of Chilean swallows (Supp. Mat. Table S1). Not even the duration of daylight, probably because of its extreme length. In contrast, mangrove swallows (16 days, 9 nests) were strongly dependent on daylight duration (Supp. Mat. Table S2), and higher daytime temperatures were associated with longer active days.

Considering the importance of the duration of the active day for daily nest attentiveness, we analyzed the timing of the first off-bout in the morning and last on-bout in the evening in an attempt to understand the variables influencing female behavior. The timing of the first daily off-bout in Ushuaia (111 first off-bouts, 16 nests) was related to calendar date: the later in the season, the earlier the females started their day, and to clutch size: larger clutches were associated with earlier off-bouts (Supp. Mat. Table S3). Mangrove swallows (29 first off-bouts, 9 nests) also tended to leave the nest earlier in the morning later in the season, and at higher temperatures (Supp. Mat. Table S4).

The timing of the last daily on-bout in Chilean swallows (112 last on-bouts, 16 nests) was not related to any of the variables analyzed (Supp. Mat. Table S5), and the same was true for mangrove swallows (23 last on-bouts, 9 nests) (Supp. Mat. Table S6).

## Discussion

Conway and Martin (2000) proposed a general framework that integrates bout duration and ambient temperature, combining embryonic heat requirements with adult energy requirements. However, few studies have fully supported this hypothesis, and our results are also far from fitting into their theoretical framework. Our results suggest that female incubation is a dynamic behavior that depends on the local ambient temperature and changes throughout the day to adequate on- and off-bout duration, trying to reduce the incubation effort whenever possible. Our results show that the optimal local temperature for the maximum bout duration rarely coincides with between off-bouts and on-bouts. Incubating females constantly adjust their behavior, which may also depend on other internal (physiology) and external processes (interspecific interactions). Our conclusions may be closer to Skutch (1962) and Austin et al. (2019), who suggested that incubating birds attend to the minimum for embryonic development, a lower limit of nest attentiveness, while the upper limit would be dictated by the physiological state of the parents. Although we suggest that the latter only happens for parents under poor condition.

Despite the differences in environmental conditions and latitude, we found common overall patterns in incubating *Tachycineta* females: 1) decreasing incubation effort with increasing ambient temperature, mainly due to a decrease in on-bout duration; 2) the positive effect of incubation day on incubation effort; 3) the positive effect of daytime on incubation effort; and 4) a positive effect of calendar date on incubation. But we also found differences that are crucial for understanding incubation behavior in these different habitats: 1) in Ushuaia, off-bouts were relatively independent of ambient temperature and became shorter as morning temperature increased, whereas in Belize they tended to become longer; 2) clutch size had a positive effect on the number and duration of bouts in females incubating in Belize, but a negative effect in Ushuaia; and 3) duration of daylight appeared to be a limiting factor only in Belize.

### Bout duration, incubation effort and ambient temperature

In both populations, females reduced their incubation efforts with increasing ambient temperature. We had expected that a reduction in incubation effort would mean a lengthening in off-bouts (Diez- Méndez et al. 2021a) but the two populations followed different strategies. Mangrove swallows did tend to lengthen their off-bouts but mainly shortened their on-bouts, while Chilean swallows achieved these numbers by shortening their on-bouts. Chilean swallows may find it challenging to lengthen their bouts given the harsh conditions. Indeed, females breeding in Ushuaia allowed their clutches to reach lower temperatures when they take as long off-bouts as the mangrove swallows, with the subsequent rewarming energy costs (Vleck 1981, Reid et al. 2002, Voss et al. 2006). This could be considered a suboptimal strategy (Ardia et al. 2009) especially with the expected low thermal inertia of small clutches (Boulton and Cassey 2012), which could explain the longer incubation periods (see below). Nonetheless, Chilean swallows also decreased in incubation effort with increasing ambient temperature, contradicting the notion of suboptimal incubation conditions at low temperatures. Conway and Martin (2000) have discussed that longer on-bouts at decreasing temperatures may be explained by females relying more on endogenous reserves or males incubating more or feeding more. In these two species, the role of males could be important, as both species are known to have males that do not incubate but can assist in slowing down egg cooling (Ospina et al. 2015, Winkler et al., unpublished), a trait that appears to be common in the Hirundinidae family (Voss et al. 2008). The observed bout duration might be explained if Chilean male swallows were more active than their conspecifics in Belize. This hypothesis needs further investigation as the currently available data is insufficient.

To add to the picture of incubation duration and attentiveness, another closely related species from the mid-latitudes of the Northern Hemisphere (tree swallows, Ardia et al. 2009, 2010) showed that off-bouts are as long as in the species examined in this study, but on-bouts are longer (estimates between 15-16 minutes). The expected nest attentiveness is consequently higher, although the authors only provided data from 24 incubation periods (∼70%). The incubation period was also shorter (∼14 days). 24h incubation appears to be similar between *Tachycineta* species, perhaps a lower limit for adequate embryo viability (Austin et al 2019), but it remains unclear whether the intermittent diurnal/continuous nocturnal incubation rate affects embryo development and incubation period duration (Diez-Méndez et al. 2021b). Overall, incubation during the active day is ∼43% higher in Ushuaia than in Hill Bank breeding females. These differences even out when nocturnal incubation is added, but Ushuaia females still require longer incubation periods. Our results show that there are different solutions to achieve similar goals depending on latitude, not only when the ambient temperature increases, but also when incubating females increase their incubation efforts during the day or later in the season.

### Clutch size and latitude

Larger clutches are favored in cold habitats, as they cool down slower during female off-bouts (egg-cooling hypothesis, Reid et al. 1999) and have a higher average temperature during on-bouts (Boulton and Cassey 2012). However larger clutches are more energetically demanding, take longer to rewarm after an off-bout (Reid et al. 2000), and require more egg movements within the clutch (Boulton and Cassey 2012). Chilean female swallows laid smaller clutches than would be expected based on their breeding latitude, but even within their clutch size range, we observed negative effects of large clutches on incubation behavior. Larger clutches tended to be associated with shorter on-bouts, possibly due to increased energy expenditure (Moreno and Sanz 1994, Thomson et al. 1998, but see de Heij et al. 2008), an increased number of daily off-bouts, and a trend towards longer active days. The latter is partly explained by an earlier first off-bout in the morning, which could be related to an earlier depletion of stored energy overnight (Bryan and Bryant 1999, Nord and Cooper 2020). For mangrove swallows, larger clutches correlated with longer off-bouts, longer on-bouts, and a trend towards fewer daily off-bouts. In this scenario, and unlike in Ushuaia, larger clutches appeared to be beneficial for female mangrove swallows as they reduced energy expenditure and rewarming episodes.

### Incubation behaviour and daily rhythms

Differences in the duration of daylight determine the daily behavior of incubating females, specifically it limits the activity of mangrove swallows. Longer daylight was associated with longer active days (and thus longer daily nest attentiveness), whereas this limitation is not as evident in females breeding in Ushuaia. Although the nocturnal hours in Belize cancel out the higher incubation effort in Ushuaia during the active day (∼7h vs. ∼10h), it is still unclear whether embryonic development during daily intermittent incubation is similar to embryonic development (Diez-Méndez et al. 2021b) or energy expenditure of the incubating female (de Heij et al. 2007, 2008) during nocturnal continuous incubation. Embryos from high latitudes may develop faster (Cooper et al. 2005) because the metabolic rate is higher during the light phase, which means shorter incubation times (Cooper et al. 2011, Austin et al. 2014). For incubating females, the shorter daylight in the tropics limits incubation behavior. Unexpectedly, we also found that higher nighttime temperatures advanced the start of the active day in mangrove swallows, resulting in longer active days on warmer days. These results could be related to the higher variability of ambient temperatures in the early morning hours (see Fig. 1) and the possibly energetic constraints of breeding females in the warm mornings.

### Duration of the incubation period

The lack of data on the total incubation period makes it difficult to adequately discuss the duration of the incubation period at different latitudes. Several studies discarded the nest attentiveness/incubation period relationship, but these approaches that analyze only the apparent incubation period or even consider hatching time as incubation (Austin et al. 2019), are usually partial (i.e., only a few days or hours of data). Patterns may only emerge when assessing the entire period (Diez-Méndez et al. 2021a). These studies also ignore the important role of partial or full incubation behavior before clutch completion (Tieleman et al. 2004), especially at low latitudes (Stoleson and Beissinger 1995, 1999). Unexpectedly, we found that Chilean swallows exhibited longer incubation periods compared to mangrove swallows, despite higher nest attentiveness and longer daylight hours, as this would promote embryonic development (Cooper et al. 2011). There are at least two non-exclusive hypotheses to explain the longer incubation periods in Chilean swallows: 1) The simplest explanation is that females at Hill Bank began incubating, either partially or fully, long before the clutch was completed, thereby shortening the apparent incubation period. This incubation behaviour has been observed in numerous passerine species (Nilsson and Svensson 1993, Ricklefs 1993, Stoleson and Beissinger 1995, Lord et al. 2011, Wang and Beissinger 2011, Podlas and Richner 2013, Diez-Méndez et al. 2020, 2021b) and specifically in other *Tachycineta* species (Ardia et al. 2006b, Ardia and Clotfelter 2007). 2) As previously mentioned, the combination of smaller clutches and low ambient temperatures may have resulted in clutches at Ushuaia being cooler during off-bouts than at Hill Bank, slowing embryonic development (Cooper et al. 2005) and lengthening the incubation periods.

In conclusion, *Tachycineta* swallows that bred in Belize and Ushuaia (Argentina) showed similar incubation constancy, but followed different strategies given the environmental conditions encountered (daylight, ambient temperature). Adapting to local conditions and reducing incubation effort whenever possible were crucial for both species. These two populations showed deviations from the latitudinal trends for life-history, which increases the importance of their assessment for understanding behavioral decisions during a critical period for later fitness values. A harsh high-latitude environment resulted in higher variability between individuals in the Chilean swallows than in the mangrove swallows, which were breeding in a more constant, mild environment with limited daylight. In this scenario, individual phenotypic plasticity in physiological and behavioral traits might be more important in Ushuaia than in Belize. Further data from the full incubation period and collecting physiological and behavioral information, including male performance, would help to fully understand latitudinal differences in incubation behavior across species and populations.

## Notes

Funding statement: The work was supported by NSF grant PIRE OISE-0730180 to DWW, CBC, and DRA. ML was supported by a fellowship from the Consejo Nacional de Investigaciones Científicas y Técnicas (CONICET). DDM was supported by a European Research Council Starting Grant BABE 805189 and is currently supported by a MSCA fellowship CZ (22_010/0008586).

### Competing Interest Statement

The authors have declared no competing interest.

