## Supplementary material for "Ambient temperature, clutch size, and daylight are the main drivers of incubation behavior in two neotropical swallows breeding 8,000 km apart"

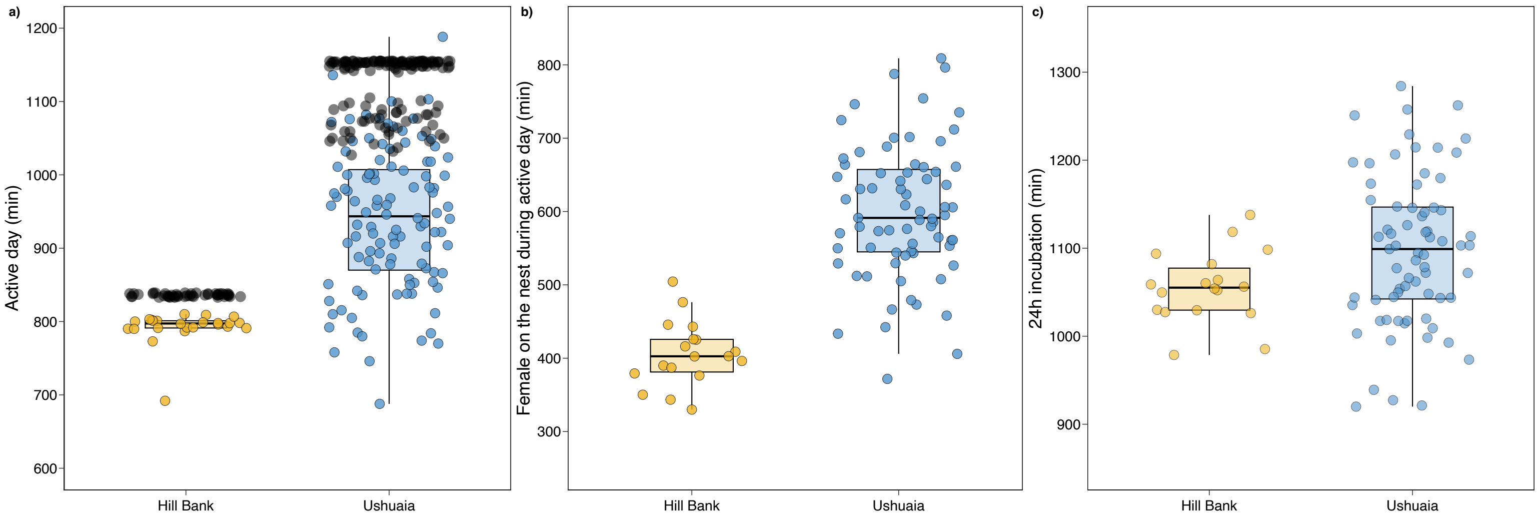


***Figure S1.*** *Raw data and derivative boxplots showing a) the duration of the active day in relation to daylight (black circles), b) sum of minutes a females spent on her nest during her active time (daily incubation), and c) 24h incubation in minutes.*

**Table S1.** Model estimates for the effect of daylight duration, clutch size, incubation day, and ambient temperature on the **duration of the active day** in **Chilean swallows**. *Nest ID* is a random intercept variable.

| Fixed effects | Estimate ± SE | z - value | P |
|---|---|---|---|
| Intercept | 944.29 ± 13.05 | 72.38 | |
| Daylight | 19.89 ± 15.9 | 1.25 | 0.211 |
| Clutch size | 27.7 ± 14.87 | 1.86 | 0.063 |
| Incubation day | -11.74 ± 10.62 | -1.11 | 0.269 |
| Temperature | 59.74 ± 89.66 | 0.67 | 0.505 |
| Temperature² | -52.4 ± 88.39 | -0.59 | 0.553 |
| **Random effects** | **Variance ± SD** | | |
| Nest | 1060 ± 32.56 | **$R^2$ fixed effects** | 0.18 |
| | | **$R^2$ model** | 0.30 |

**Table S2.** Model estimates for the effect of daylight duration, clutch size, incubation day, and ambient temperature on the **duration of the active day** in **mangrove swallows**. *Nest ID* is a random intercept variable. Significant results are highlighted in bold.

| Fixed effects | Estimate ± SE | z - value | P |
|---|---|---|---|
| Intercept | 795.05 ± 1.961 | 405.5 | |
| **Daylight** | **6.438 ± 1.536** | **4.2** | **<0.001** |
| Clutch size | -3.183 ± 2.019 | -1.6 | 0.115 |
| Incubation day | 0.437 ± 1.455 | 0.3 | 0.764 |
| **Temperature** | **10.49 ± 4.373** | **2.4** | **0.016** |
| Temperature² | -0.56 ± 4.263 | -0.1 | 0.896 |
| **Random effects** | **Variance ± SD** | | |
| Nest | 1060 ± 32.56 | **$R^2$ fixed effects** | 0.62 |
| | | **$R^2$ model** | 0.91 |

**Table S3.** Model estimates for the effect of calendar date, clutch size, incubation day, and ambient temperature on the **timing of the first daily off-bout** in **Chilean swallows**. *Nest ID* is a random intercept variable. Significant results are highlighted in bold.

| Fixed effects | Estimate ± SE | z - value | P |
|---|---|---|---|
| Intercept | 115.78 ± 9.020 | 12.84 | |
| **Date** | **-24.74 ± 9.892** | **-2.5** | **0.012** |
| **Clutch size** | **-25.03 ± 9.681** | **-2.59** | **0.010** |
| Incubation day | 4.84 ± 6.298 | 0.77 | 0.442 |
| Temperature | -25.43 ± 62.969 | -0.40 | 0.686 |
| $Temperature^2$ | -75.45 ± 62.906 | -1.20 | 0.230 |
| **Random effects** | **Variance ± SD** | | |
| Nest | 609.8 ± 24.69 | **$R^2$ fixed effects** | 0.18 |
| | | **$R^2$ model** | 0.29 |

**Table S4.** Model estimates for the effect of calendar date, clutch size, incubation day, and ambient temperature on the **timing of the first daily off-bout** in **mangrove swallows**. *Nest ID* is a random intercept variable. Significant results are highlighted in bold.

| Fixed effects | Estimate ± SE | z - value | P |
|---|---|---|---|
| Intercept | 12.38 ± 0.852 | 14.53 | |
| Date | -2.23 ± 1.264 | -1.76 | 0.078 |
| Clutch size | 1.00 ± 0.968 | 1.03 | 0.303 |
| Incubation day | -0.56 ± 1.040 | -0.54 | 0.592 |
| **Temperature** | **-13.67 ± 6.776** | **-2.02** | **0.044** |
| $Temperature^2$ | 4.11 ± 5.175 | 0.79 | 0.427 |
| **Random effects** | **Variance ± SD** | | |
| Nest | 0.00 ± 0.0001 | **$R^2$ fixed effects** | 0.19 |
| | | **$R^2$ model** | 0.19 |

**Table S5.** Model estimates for the effect of calendar date, clutch size, incubation day, and ambient temperature on the **timing of the last daily on-bout** in **Chilean swallows**. *Nest ID* is a random intercept variable.

| **Fixed effects** | **Estimate ± SE** | **z - value** | **P** |
|---|---|---|---|
| Intercept | 66.82 ± 5.519 | 12.11 | |
| Date | 4.25 ± 5.914 | 0.72 | 0.473 |
| Clutch size | -8.96 ± 6.002 | -1.49 | 0.135 |
| Incubation day | 2.55 ± 3.677 | 0.69 | 0.488 |
| Temperature | -48.45 ± 38.190 | -1.27 | 0.205 |
| Temperature$^2$ | 37.09 ± 37.042 | 1.00 | 0.317 |
| **Random effects** | **Variance ± SD** | | |
| Nest | 241.8 ± 15.55 | **$R^2$ fixed effects** | 0.09 |
| | | **$R^2$ model** | 0.24 |

**Table S6.** Model estimates for the effect of calendar date, clutch size, incubation day, and ambient temperature on the **timing of the last daily on-bout** in **mangrove swallows**. *Nest ID* is a random intercept variable.

| **Fixed effects** | **Estimate ± SE** | **z - value** | **P** |
|---|---|---|---|
| Intercept | 28.45 ± 0.861 | 33.04 | |
| Date | -1.48 ± 0.980 | -1.51 | 0.131 |
| Clutch size | 1.50 ± 0.915 | 1.63 | 0.102 |
| Incubation day | 0.36 ± 1.053 | 0.34 | 0.732 |
| Temperature | -3.42 ± 4.318 | -0.79 | 0.429 |
| Temperature$^2$ | 4.22 ± 4.499 | 0.94 | 0.348 |
| **Random effects** | **Variance ± SD** | | |
| Nest | 0.00 ± 0.0002 | **$R^2$ fixed effects** | 0.32 |
| | | **$R^2$ model** | 0.32 |
